# Adaptation with naturalistic textures in macaque V1 and V2

**DOI:** 10.1101/2023.12.05.569995

**Authors:** Aida Davila, Adam Kohn

## Abstract

Adaptation affects neuronal responsivity and selectivity throughout the visual hierarchy. However, because most prior studies have tailored stimuli to a single brain area of interest, we have a poor understanding of how exposure to a particular image alters responsivity and tuning at different stages of visual processing. Here we assess how adaptation with naturalistic textures alters neuronal responsivity and selectivity in primary visual cortex (V1) and area V2 of macaque monkeys. Neurons in both areas respond to textures, but V2 neurons are sensitive to high-order statistics which do not strongly modulate V1 responsivity. We tested the specificity of adaptation in each area with textures and spectrally-matched ‘noise’ stimuli. Adaptation reduced responsivity in both V1 and V2, but only in V2 was the reduction dependent on the presence of higher-order texture statistics. Despite this specificity, the texture information provided by single neurons and populations was reduced after adaptation, in both V1 and V2. Our results suggest that adaptation effects for a given feature are induced at the stage of processing that tuning for those features first arises and that stimulus-specific adaptation effects need not result in improved sensory coding.

**Significance statement:** Nearly all sensory neurons adapt to recent input. However, how the adjustment triggered by a particular input is distributed across brain areas and how these changes contribute to sensory processing are poorly understood. Here we explore adaptation with naturalistic textures, for which neurons in primary visual cortex and area V2 have differing selectivity. Adaptation reduced responsivity in both areas, but the effects in V2 alone depended on the presence of the higher-level texture statistics to which V2 neurons are sensitive. Though prior work with simpler stimuli has argued that stimulus-specific adaptation improves visual encoding, we found adaptation reduced texture information. Thus, adaptation need not improve visual encoding, suggesting its effects may serve some other purpose.

---

Our immediate previous sensory experience—adaptation—affects neuronal responses to current stimuli, allowing our sensory systems to adjust to the environment (Kohn, 2007; Rieke and Rudd, 2009; Webster, 2015; Weber and Fairhall, 2019). In the visual system, adaptation has been found to affect neuronal responses in the retina (Brown and Masland, 2001; Demb, 2008), LGN (Solomon et al., 2004; Camp et al., 2009), V1 (Movshon and Lennie, 1979; Ohzawa et al., 1982; Dragoi et al., 2002; Ghisovan et al., 2009; Patterson et al., 2013), V2 (Crowder et al., 2006), V4 (Tolias et al., 2005), MT (Priebe et al., 2002; Kohn and Movshon, 2003), IT (Sawamura et al., 2006; Fabbrini et al., 2019), and other visual areas.

Most work on neuronal adaptation has used stimuli tailored to the area being investigated. Because different areas are studied with different stimuli, it remains unclear how adaptation with a given image affects encoding that is distributed across stages of the visual hierarchy (Solomon and Kohn, 2014). For a complex stimulus for which different image attributes are encoded at different stages of processing, how are the effects induced by adaptation distributed across those areas? Do higher visual areas adapt to the stimulus features they encode, or are the effects in those areas mostly inherited from earlier visual areas as previous work has often described (Kohn and Movshon, 2003; Dhruv and Carandini, 2014; Patterson et al., 2014)? Additionally, do changes in neuronal responsivity or selectivity after adaptation, whether locally generated or inherited, improve visual encoding? Improved encoding has been observed in primary sensory cortex but it is not clear whether the encoding of more complex visual statistics in higher cortex is improved after adaptation (Müller et al., 1999; Sharpee et al., 2006; Gutnisky and Dragoi, 2008; Ghodrati et al., 2019; Jin et al., 2019).

Here we assess the specificity and consequences of adaptation with naturalistic textures in primary visual cortex (V1) and area V2 of the macaque monkey. Naturalistic textures are composed of homogenous, repeating structures which can be summarized by a set of summary statistics, consisting of both low- and high-level features (Portilla and Simoncelli, 2000). Low-level features can be described by the output of Gabor-like filters which capture luminance, contrast, and spectral information. High-level features consist of the correlation of filter outputs across space and spatial scales, which produce borders and structures within textures. V1 neurons are insensitive to high-level features of a texture, responding similarly to naturalistic textures and to spectrally-matched noise images (Freeman et al., 2013; Ziemba et al., 2016). In contrast, V2 neurons respond more strongly to textures which contain higher-level features than those which do not (Freeman et al., 2013; Ziemba et al., 2016, 2019).

We aimed to understand 1) whether V1 and V2 neurons adapt differently to texture and spectrally-matched noise stimuli, reflecting the features of those stimuli for which they are selective; and 2) how changes in neuronal responsivity after adaptation affect the encoding of textures. We find that brief exposure to a texture or spectrally matched noise image reduces responsivity in both V1 and V2 units. The adaptation effects in V2 but not V1 display specificity for texture statistics. This specificity is evident in comparisons between adaptation effects measured with texture and noise images and in comparisons of effects across different textures. In both areas, adaptation reduced the information about textures provided by the responses of single neurons and neuronal populations.

## Materials and methods

### Surgery

Recordings were conducted in four adult male monkeys (*Macaca fascicularis*). Prior to the induction of anesthesia with ketamine (10 mg/kg) animals were administered glycopyrrolate (0.01 mg/kg) and diazepam (1.5 mg/kg). Once anesthesia was confirmed, animals were intubated and placed on 1.0%-2.5% isoflurane in a 98% O_2_-2% CO_2_ mixture. Intravenous catheters were then inserted into the saphenous veins of both legs. Animals were placed in a stereotaxic device where a craniotomy and durotomy was performed over V1 (centered ∼5 mm posterior to the lunate sulcus and ∼10 mm lateral to the midline). Postsurgical anesthesia was preserved with a venous infusion of sufentanil citrate (6–24 microg/kg/h, adjusted as needed) in Normosol solution with dextrose. To minimize eye movements vecuronium bromide (150 microg/kg/h) was also administered via venous infusion. Vital signs, comprising of heart rate, SpO2, ECG, blood pressure, EEG, end-tidal CO2, airway pressure, core temperature, and urinary output and specific gravity, were continuously monitored throughout the recordings to ensure suitable anesthesia and physiological state. Rectal temperature was maintained near 37°C using heating pads. Pupils were dilated with topical atropine. Gas-permeable contact lenses were used to protect the corneas. Additional supplementary lenses were used to bring the retinal image into focus. Antibiotics (Ceftliflex, 2.2 mg/kg) and a corticosteroid (dexamethasone, 1 mg/kg) were administered daily. All procedures were approved by the Institutional Animal Care and Use Committee of the Albert Einstein College of Medicine and were in compliance with the guidelines set forth in the National Institutes of Health’s *Guide for the Care and Use of Laboratory Animals*.

### Recordings

Neuropixel Phase 3B probes placed on a custom 3D-printed probe holder were lowered into the cortex using a microelectrode drive (Alpha Omega). Each recording session involved 2-4 Neuropixel probes, arranged mediolaterally. Neuropixel probes were sharpened prior to insertion. No guide tubes were used during insertion. Once Neuropixel probes were positioned in the brain, the craniotomy was filled with agar or Dura-Gel to prevent desiccation of the cortex.

In one animal, neural responses were acquired from the 384 channels extending over a distance of 3.84 mm, closest to the probe tip which was in V2. In the remaining 3 animals we used a probe configuration which recorded from 384 channels in a staggered manner, spanning 7.68 mm of the probe and all layers of V1 and V2. Data were acquired with SpikeGLX or OpenEphys software.

Raw data were spike sorted with Kilosort 2.5, which clusters units based on waveform shape (Pachitariu et al., 2016). Kilosort output was then manually curated using Phy, which involved merging and splitting units and confirming waveform templates were from a single neuron.

### Visual stimuli

Stimuli were presented on a calibrated cathode ray tube monitor (Hewlett Packard p1230; 1024 × 768 pixels; 100 Hz frame rate, ∼40 cd/m^2^ mean luminance), using custom OpenGL software (EXPO; P. Lennie). The monitor was placed approximately 90-110 cm from the animal. To determine stimulus placement, we measured the spatial receptive field of each recording site by presenting small drifting gratings at 169 different screen locations, spanning roughly a 5° × 5° region of visual space. Receptive field data was collected for each eye independently. Recorded voltage traces were high pass filtered at 300 Hz and thresholded, and V1 and V2 aggregate receptive fields were computed from these multiunit responses. Subsequent stimuli were centered to drive as many units as possible in both areas. Units were only included in our analysis if their receptive fields were entirely covered by the 8° stimulus.

We used textures synthesized with the Portilla and Simoncelli algorithm (Portilla and Simoncelli, 2000). The algorithm quantifies a variety of statistics of an input texture and synthesizes a new image enforcing the measured statistics. Texture synthesis for each image was initialized with a Gaussian white noise image. Analyzed statistics from the Portilla and Simoncelli algorithm were then iteratively applied (n=50) so that the noise image resembled the original texture. Textures were synthesized with 4 scales and orientations of Gabor-like filters, and correlations were derived within a 15-pixel square. Input textures were obtained from the Brodatz texture database (Abdelmounaime and Dong-Chen, 2013), Kylberg database (Kylberg, 2011), Oulu texture database (Lee and Bang, 2019), and Salzburg texture images (https://www.wavelab.at/sources/STex/). Each input texture (256 x 256 pixels) was gray scaled and served as the basis for a texture “family”. A total of 23 texture families were used. For each texture family the noise counterpart was synthesized by randomizing the Fourier phase of the synthesized texture and then performing an inverse Fourier transform. All synthesized textures and matched noise images were adjusted to have the same contrast (RMS=0.314) and mean luminance (128, on an 8-bit range). Textures and noise counterparts were synthesized to be 512 × 512 pixels. For all experiments we presented stimuli windowed with a 8° aperture (40-54 pixels/degree), centering the aperture at random locations of the synthesized texture or noise stimuli, while ensuring that the aperture border did not extend past the edge of image. Each randomized centering of a texture family thus contained the same statistics but differed at a pixel level. We refer to each randomized centering as a sample of a texture family.

Each trial began with the presentation of an adapter which could be a uniform gray background (control), texture or counterpart noise image, followed by an inter-stimulus interval and a test stimulus. For texture and noise adapters, we refreshed the sample of the stimulus every 100 ms, so that adaptation was defined by the statistics of the image and not the pixel-level content. Similarly, the test stimulus consisted of three unique 100 ms samples of a texture or its noise counterpart, presented in continuous succession. The ensemble of test stimuli consisted of five textures, five counterpart noise images, and a blank. Importantly, one test stimulus was from the same texture family as the adapter and the rest were from a different family. This stimulus ensemble thus allowed us to test adaptation specificity both for textures compared to their noise counterparts, as well as for specificity among different texture families. Each adaptation condition was presented 76-129 times (mean=100). Stimuli were presented either binocularly or monocularly.

### Neural data analysis

Data analysis was performed in MATLAB (MathWorks). Each neuron’s response was measured as the spike count in a 300 ms window, beginning 70 ms after stimulus onset and ending 70 ms after stimulus offset to account for the response latency. Pre-adaptation spontaneous activity was calculated as the spike count during the blank test stimulus which followed a blank adapter; post-adaptation spontaneous activity was defined similarly, after the presentation of the texture or noise adapter. Units included in our analysis had to have a mean response higher than the mean spontaneous activity plus 1.5 times the standard deviation of that response to at least one of the 10 test stimuli (5 different textures and 5 noise stimuli). In addition, neurons were required to have a median spike count of at least 1 spike/trial. For each neuron, adaptation indices, modulation indices, and auROC values were calculated only for textures that evoked responses that exceeded these criteria, either before or after adaptation.

We calculated a modulation index for each unit and texture family by taking the difference in evoked responses between textures and their noise counterpart and dividing by their sum. The modulation index was measured using responses to test stimuli presented after a blank adapter. A modulation index was calculated separately for each texture family and each neuron.

To quantify adaptation, we calculated an adaptation index from the evoked responses, by subtracting the unadapted response from the adapted response, and dividing by their sum. To quantify single unit texture information, we computed the discriminability between texture families afforded by the measured responses, using the area under the receiver operating characteristic curve (auROC). The auROC was computed for the four possible pairs of texture families, in which one of the textures was always the adapted texture and the other was one of the four other textures. We measured discriminability in all cases for which the response to at least one of the two textures met the inclusion criteria, in either the adapted or un-adapted condition. For all plotted auROC values, we used the absolute values of the deviation from chance performance (a value of 0.5). Trials were matched across conditions for all auROC analyses.

We used bootstrap analysis to evaluate the significance of selectivity for textures compared to noise images (the modulation index), for selectivity for different texture families (auROC), and for the strength of adaptation (adaptation index). Bootstraps for modulation index values were performed by creating 1,000 ‘texture’ and ‘spectral noise’ data sets by randomly choosing subsets of responses for either texture or noise responses with replacement. We then calculated the modulation index for each of the 1,000 responses and evaluated significance by indexing the values observed in the data in this null distribution. The statistical significance of the auROC and adaptation index values were calculated similarly.

To quantify texture information in neuronal populations we used linear discriminant analysis (LDA) with 10-fold cross validation. We performed a 2-way discrimination task between texture families, in which one texture was always matched to the adapter and the other was one of the remaining textures (four possible pairs). Firing rates were normalized (z-scored) prior to training the classifier. Classifiers were trained for adapted and un-adapted responses separately, using the same units and number of trials per condition. Using regularization to reduce over-fitting, we also tested LDA performance as a function of population size. Reported classifier performance values are on held out data.

## Results

We used Neuropixel probes to record V1 (n=329) and V2 (n=435) neurons in 4 anesthetized monkeys (Fig. 1A). In each recording session, we measured the spatial receptive field of each recorded unit (see Methods). The stimulus location was then chosen to cover the aggregate receptive fields of the sampled V1 and V2 units (Fig. 1B, plotted in green and blue, respectively).

**Fig. 1.**
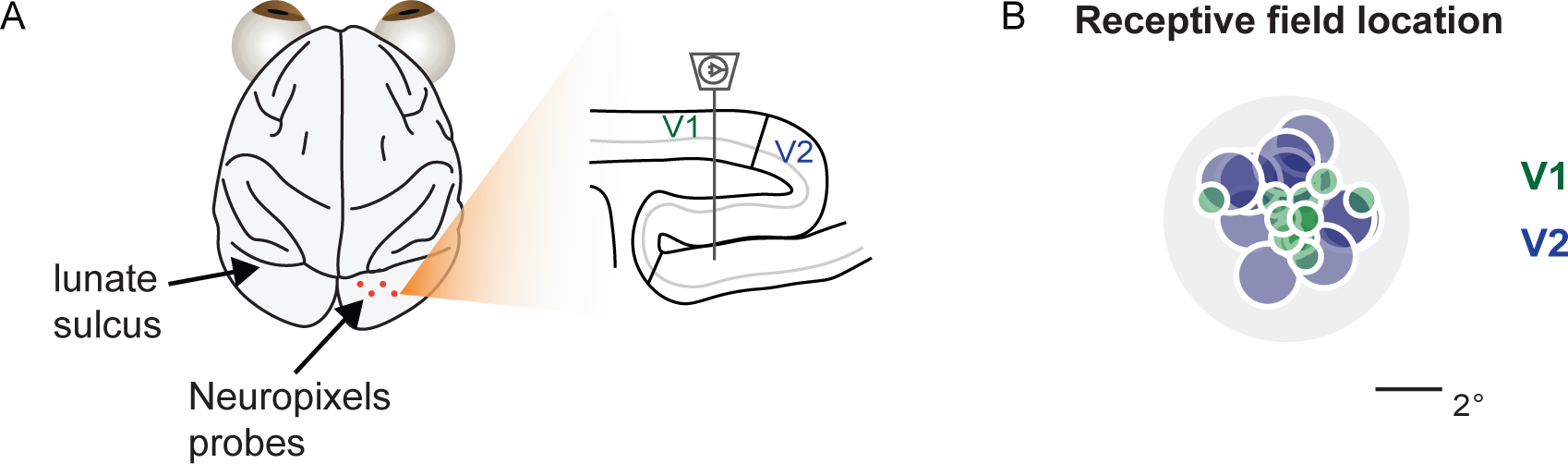
Recording approach. **(A)** Neuropixels probes (position on cortical surface indicated in orange) were lowered to record V1 and V2 units simultaneously, indicated in sagittal view (right). **(B)** Aggregate receptive field locations for V1 (green) and V2 (blue) units, across all sessions, with respect to the center of the 8° stimulus (gray).

We synthesized stimuli from different texture images, using the Portilla and Simoncelli algorithm (Portilla and Simoncelli, 2000). Each texture image was used to generate a texture family, or a set of synthetic textures with the same summary statistics. To test whether units adapted to the higher- or lower-order (spectral) statistics of a texture family, we also synthesized control images for each texture family that were matched for spectral content but not other statistics (termed hereafter, ‘noise’ images). All stimuli were matched in contrast and mean

On each trial, we presented a 600 ms adapter followed by a 100 ms inter-stimulus interval and a 300 ms test stimulus (Fig. 2, left). The adapter was either a texture, its counterpart noise image or a blank screen (Fig. 2, middle). For texture and noise adapters, we refreshed the sample of the stimulus every 100 ms, so that adaptation was defined by the statistics of the image and not the pixel-level content. Similarly, the test stimulus consisted of three unique 100 ms samples of a texture or its noise counterpart, presented in continuous succession. The ensemble of test stimuli consisted of five textures, five counterpart noise images, and a blank. Importantly, one test stimulus was from the same texture family as the adapter (Fig. 2, right, orange) and the others from a different family (Fig. 2, right, yellow). This stimulus ensemble thus allowed us to test adaptation specificity both for textures compared to their noise counterparts, as well as for specificity among different texture families.

**Fig. 2.**
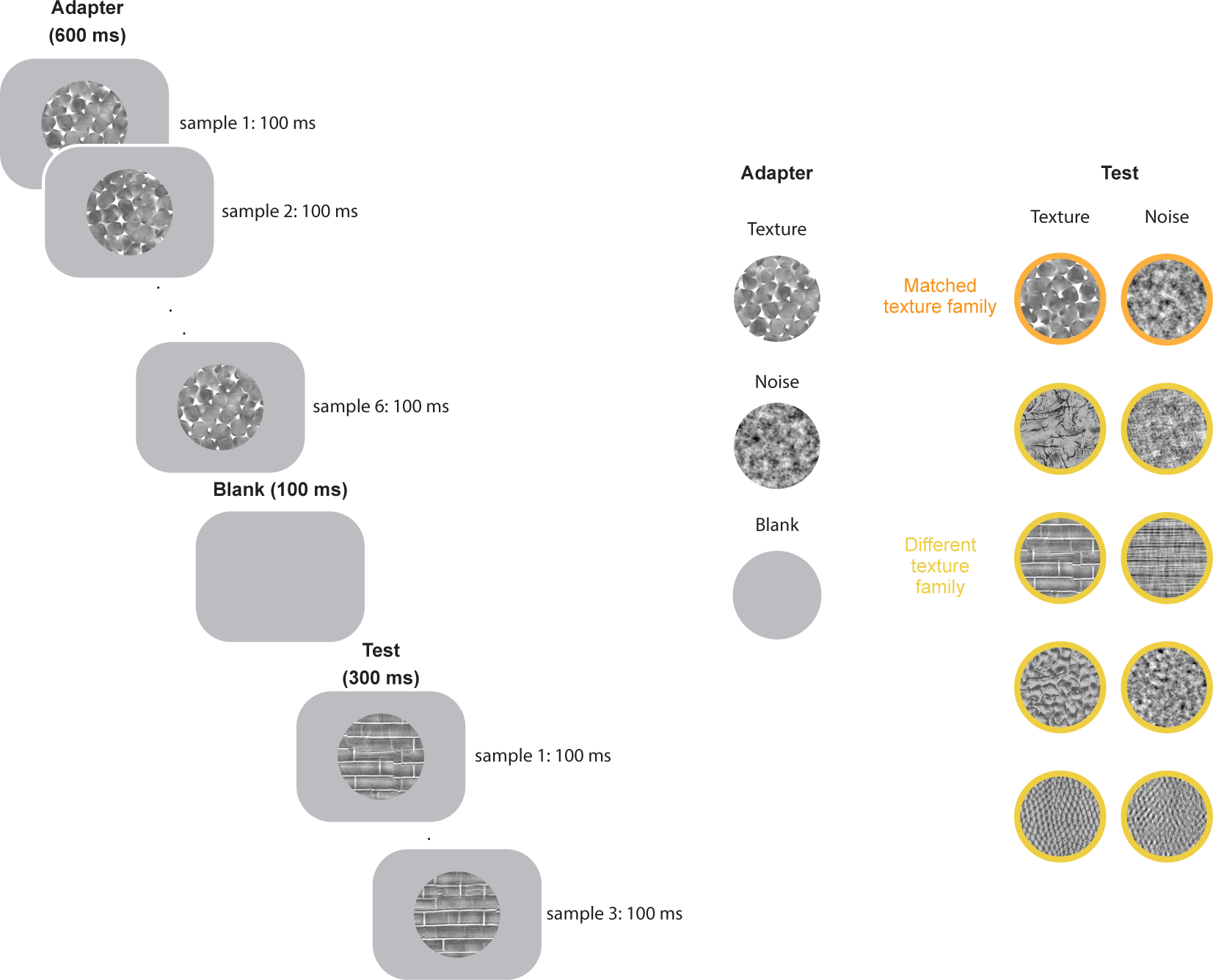
Experimental design. *Left*: Each trial consisted of an adapter (600 ms) followed by a blank (100 ms) and a test stimulus (300 ms) *Middle*: We used 3 different types of adapters. *Right*: Test stimuli consisted of 5 different texture families, 5 corresponding noise images, or a blank stimulus. One of the test stimuli was always from the same texture family as the adapter, though different samples. This design allowed us to test adaptation effects within a texture family (right, orange) or across texture families (right, yellow).

### Sensitivity for texture statistics

We first assessed the sensitivity of each cell for texture statistics, by calculating a modulation index using the responses to test stimuli that followed a blank adapter (i.e., control responses). The modulation index was defined as the difference between responses to synthesized textures and their noise counterpart normalized by their sum (Freeman et al., 2013; Ziemba et al., 2016).

Modulation indices for both V1 and V2 units extended over a substantial range, with the responses of some units being quite similar for textures and noise stimuli (a value near 0, example units in Fig. 3A, left) and some showing a greater difference between the two types of stimuli (a value greater than 0, example units in Fig. 3A, right). Consistent with previous studies (Freeman et al., 2013; Ziemba et al., 2016, 2019), the mean modulation index was higher in V2 than V1 (0.08±0.006 vs. 0.03±0.007, p<0.001, unpaired t-test; Fig. 3B). We also determined the percentage of cases for which the modulation index was statistically different from 0 (using a 95% confidence interval from a bootstrap analysis; Fig. 3B, filled bars). There was a slightly higher percentage of significant cases in V2 (44%) than V1 (36%), and these more often had positive modulation indices, indicating stronger responses to textures than noise. To understand how the sensitivity to texture statistics arose in time, we calculate population peri-stimulus histograms from the normalized responses of each cell. On average differences between responses to textures and noise arose briefly after response onset and were greater in magnitude in V2 (Fig. 3C, bottom) than V1 (Fig. 3C, top).

**Fig. 3.**
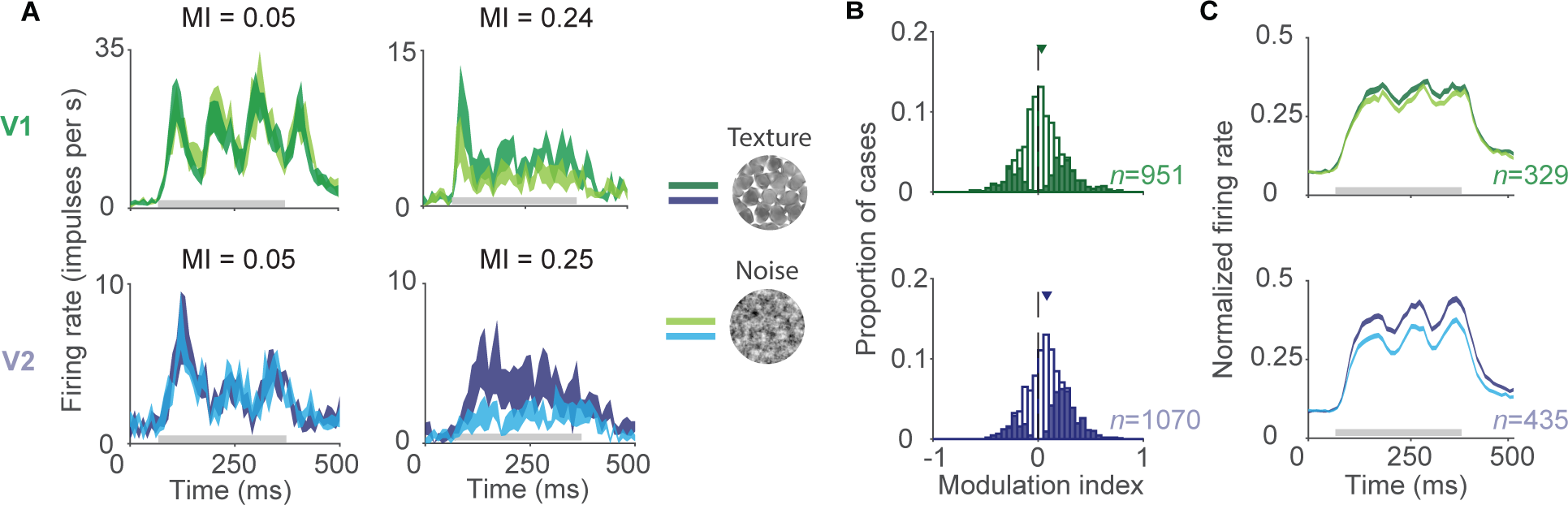
Sensitivity for texture statistics. **(A)** Example responses of V1 (top) and V2 (bottom) units to naturalistic textures (darker traces) and their noise counterparts (light traces). Thickness of line depicts S.E.M. across texture families per unit. Units in both V1 and V2 could show high (left) or low (right) modulation between the two stimulus classes. Gray line indicates the time during which the stimulus was presented. **(B)** Modulation indices for V1 (top) and V2 (bottom) units. Shaded bars represent significant cases. **(C)** Population average PSTHs for V1 (top) and V2 (bottom) units.

We noted that the distribution of V2 modulation indices was less skewed towards positive values than those reported in previous studies (Freeman et al., 2013; Ziemba et al., 2016). These studies found selectivity for texture statistics in V2 arose in part from the receptive field surround (Ziemba et al., 2018), so that larger naturalistic textures were associated with greater modulation indices. We thought that the difference between our values and theirs might thus be due to differences in the stimulus’ centering of our recorded V2 units (Fig. 1B, blue circles), which could affect how the surround was engaged. To evaluate this possibility, we analyzed a separate data set in which stimuli were well-centered on the sampled V2 units (n=237). In these data, V2 units had modulation indices skewed towards more positive values (0.16±0.006 vs. 0.08±0.006, p<0.0001 unpaired t-test), similar to Freeman et al. (2016).

### Adaptation specificity for textures vs. noise images

We next tested whether the different sensitivity to texture statistics in V1 and V2 would lead to different adaptation specificity in the two areas. Since V1 neurons do not distinguish strongly between texture and noise stimuli, one might expect adaptation effects to be similar in magnitude for texture or noise adapters, when measured with either texture or noise test stimuli (Fig. 4). That is, since textures and noise are indistinguishable for many V1 cells, we would expect no adaptation specificity, based on the adapter and test stimulus type. In contrast, many V2 neurons do distinguish between textures and noise, so one might expect that adaptation effects in V2 would be stronger when adapter and test stimuli are matched in statistics than when they differ (e.g., Fig. 4A vs Fig. 4B). In addition, we might expect that texture adapters would induce stronger effects since they are more potent than noise images in V2 (Fig. 4A, B vs Fig. 4C, D).

**Fig. 4.**
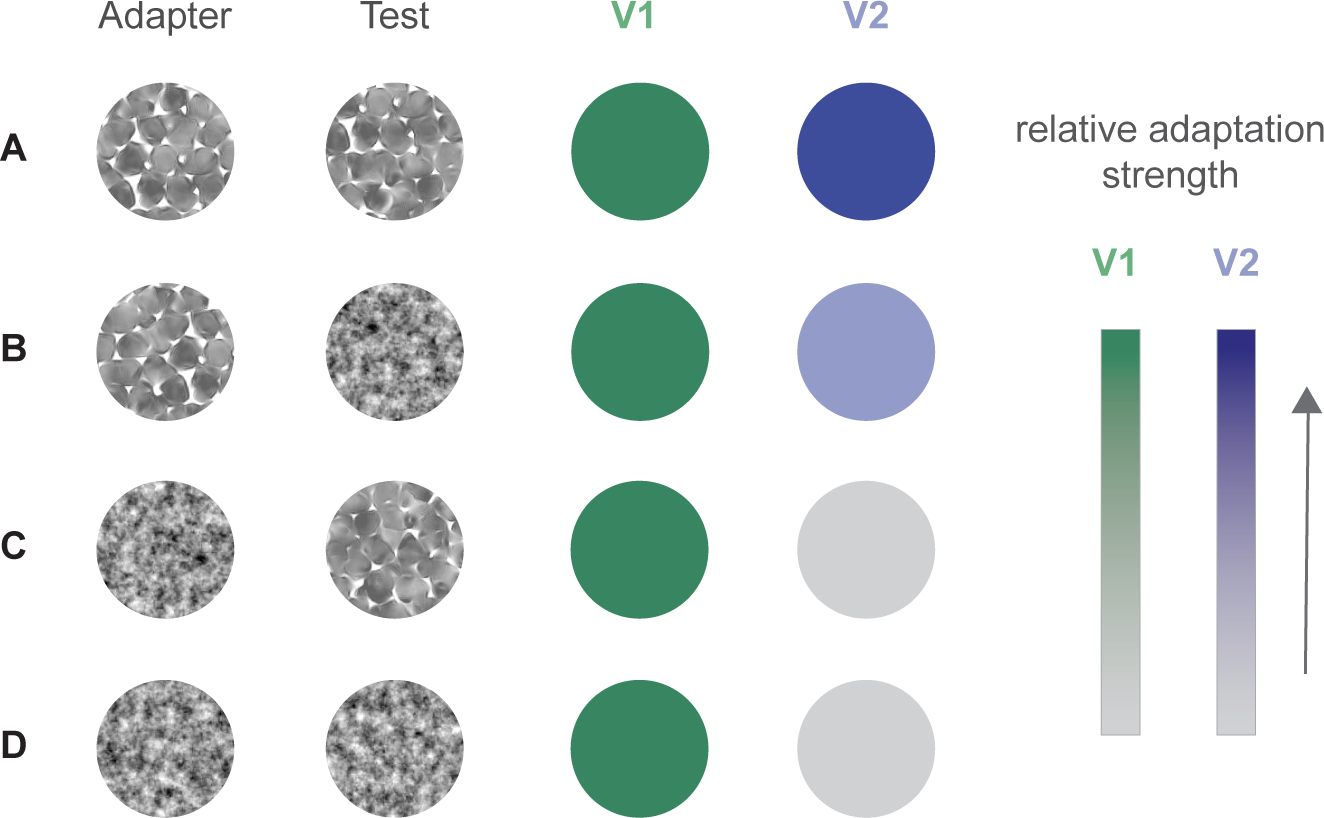
Predictions of adaptation effects for V1 and V2. **(A)** Adapting to a texture and testing to a texture should produce strong adaptation effects in V1 (green) and V2 (blue). **(B)** Adapting to a texture and testing to noise should elicit strong adaptation effects in V1 (green) but not V2 (light blue). **(C)** Adaptation to noise and testing with a texture should produce strong adaptation effects in V1 (green) but not V2 (gray). **(D)** Adapting with noise and testing with noise should produce strong adaptation effects in V1 (green) but not V2 (gray).

We measured how strongly responsivity was altered by adaptation using an adaptation index, defined as the difference in responses between unadapted and adapted conditions divided by their sum. Adaptation effects could manifest as either response facilitation (adaptation index less than zero) or suppression (greater than zero).

We first examined how texture adapters affected responsivity. Texture adapters weakened V1 responses for both texture (0.07±0.01, p<0.001, t-test; Fig. 5A, upright) and noise test stimuli (0.08±0.01, p<0.001, t-test; Fig. 5A, flipped), and the effects for these two types of test stimuli were similar in magnitude (p=0.9, unpaired t-test). V2 units also responded less strongly to texture (0.07 ±0.01, p<0.001, t-test; Fig. 5B, upright) and noise test stimuli (0.03±0.02; p=0.02, unpaired t-test; Fig. 5B, flipped) following a texture adapter. Unlike V1, however, V2 responses to texture test stimuli were reduced more than to noise stimuli (p=0.05, unpaired t-test). Thus, in V2 but not V1 the effects induced by texture adapters depended on the presence of higher-level texture statistics in the test stimulus.

**Figure 5.**
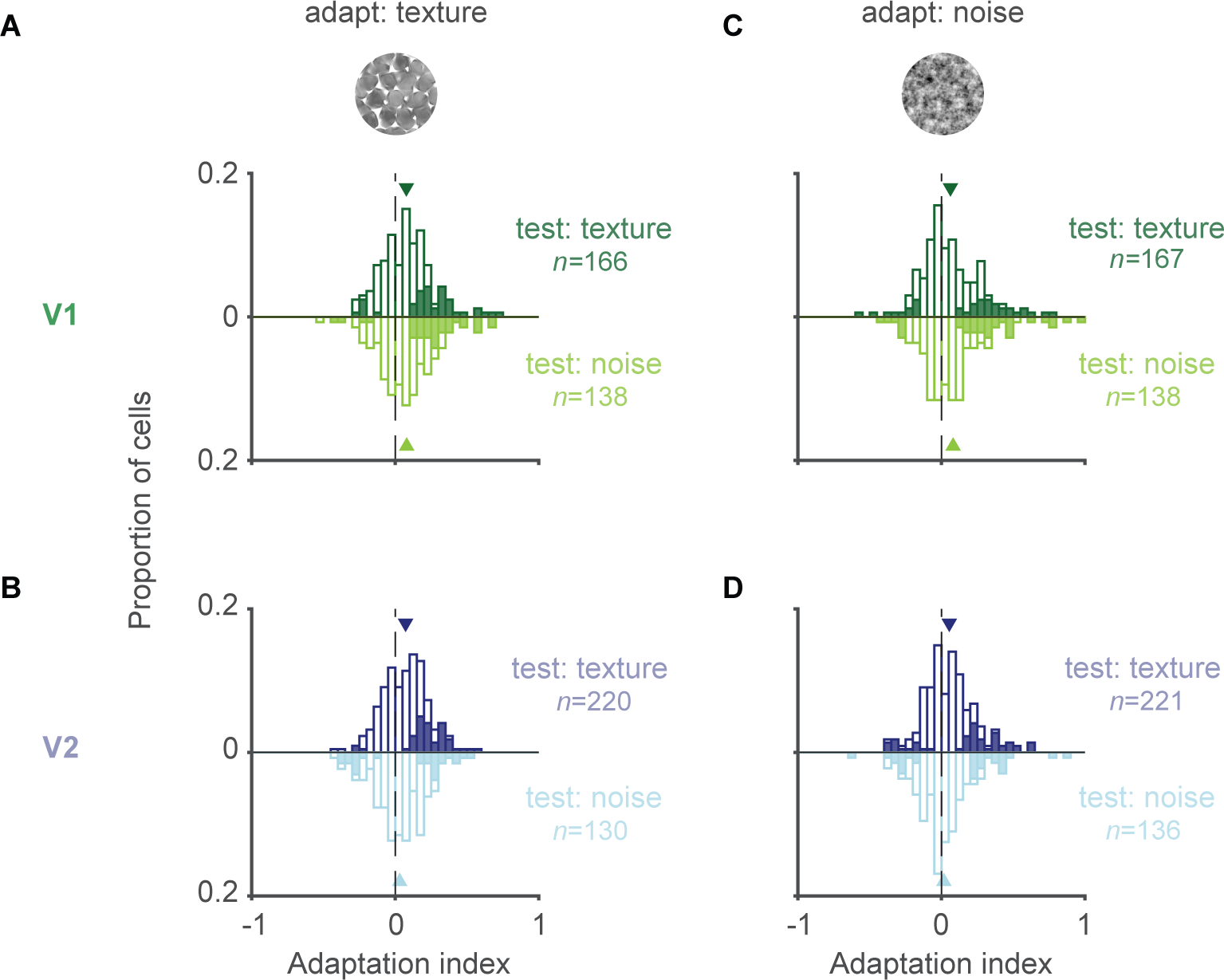
Dependence of adaptation effects on higher order texture statistics in V1 and V2. **(A, B)** Distribution of adaptation indices for V1(A) and V2 (B) after adapting with a texture and testing with either a texture (upright, dark traces) or noise (flipped, light traces) stimulus. Cases with significant adaptation indices are shown with filled bars. Mean of distribution is indicated by triangle. **(C, D)** Distribution of adaptation indices for V1 (C) and V2 (D) after adapting with a noise stimulus and testing with either a texture (upright, dark traces) or noise (flipped, light traces) stimulus.

Next, we tested whether adapters which lacked higher-level statistics (noise images) induced weaker or stronger changes in responsivity than texture adapters. Noise adapters reduced V1 responsivity for both texture (0.06±0.01, p<0.001, t-test; Fig. 5C, upright) and noise test stimuli (0.08±0.02, p<0.001, t-test; Fig. 5C, flipped). This response reduction in V1 units was indistinguishable for the two test stimuli (p=0.47, unpaired t-test; Fig. 5C). Further, the effects induced by texture and noise adapters were similar both for texture (p=0.48, unpaired t-test) and noise test stimuli (p=0.97, unpaired t-test).

In V2, noise adapters reduced responses to texture test stimuli (0.05±0.01, p<0.001, unpaired t-test; Fig. 5D, upright) and more weakly affected responses to noise stimuli (0.02±0.02, p=0.36, unpaired t-test; Fig. 5D, flipped), though the adaptation indices for the two test stimuli was not significantly different (p=0.07, unmatched t-test; Fig. 5D). Although V2 units had slightly stronger adaptation indices for texture than noise adapters, the differences were small for both texture (p=0.27, unpaired t-test) and noise (p=0.55, unpaired t-test) test stimuli. Thus, the statistics of the test stimulus were more important than those of the adapter for revealing adaptation specificity.

The recorded units showed a wide range of sensitivities to texture statistics: some cells responded equally to textures and noise images, others responded more strongly to one or the other. We wondered whether adaptation specificity (or its absence) was related to each unit’s sensitivity to texture statistics. To evaluate this possibility, we focused on units in V1 (n=62) and V2 (n=134) that had a significant modulation index for the texture used as the adapter. If adaptation specificity for texture statistics were dependent on single neuron selectivity, adaptation specificity in V1 and V2 should be higher in units with a significant modulation index. In V1, this was not the case: units with a significant modulation index had similar adaptation indices for textures and noise test stimuli after adaptation with a texture (0.05±0.02 vs. 0.06±0.03, p=0.7, unpaired t-test) or noise (0.04±0.02 vs. 0.05±0.02, p=0.9, unpaired t-test). Even in those V1 units with a significant positive modulation index, we found no evidence of specificity (n=44; for texture adapters, the adaptation indices for test textures and noise images were 0.08±0.02 vs. 0.05±0.03, respectively, paired t-test p=0.46). In V2, units with a significant modulation index had higher adaptation indices for texture compared to noise test stimuli following a texture adapter (0.08±0.01 vs. 0±0.02, respectively, p<0.0001, unpaired t-test) or noise adapter (0.06±0.01 vs. 0±0.02, p=0.02, unpaired t-test).

Thus, although we observe differences in adaptation strength in V2 for texture and noise test stimuli after a texture adapter, this specificity was not obviously related to each unit’s modulation index but rather seems to reflect the sensitivity of the V2 network to texture statistics. In V1, there is little difference between the effects induced or measured with either textures or noise images, even in units with significant modulation indices.

### Specificity of adaptation effects between texture families

We next asked whether adaptation altered responses solely for the adapted texture family, or whether responses to other texture families were also affected.

We calculated adaptation indices for each of the 4 test texture families that differed from the adapter and compared those indices to those measured for test stimuli matched to the adapter. V1 responsivity was reduced after adaptation both for test textures matched to (0.07±0.01, p<0.001, t-test; Fig. 6A, upright) and different from (0.06±0.01, p<0.001, t-test; Fig. 6A, flipped) the texture adapter. The adaptation indices were statistically indistinguishable for the two cases (p=0.32, unpaired t-test for difference). V2 units also adapted both to test textures matched to (0.07±0.01, p<0.001, t-test; Fig. 6B, upright) and different from (0.04 ± 0.01, p<0.001, t-test; Fig. 6B, flipped) the texture adapter. However, in V2 the adaptation indices were larger for test textures matched to the adapter (p=0.02, unpaired t-test).

**Fig. 6.**
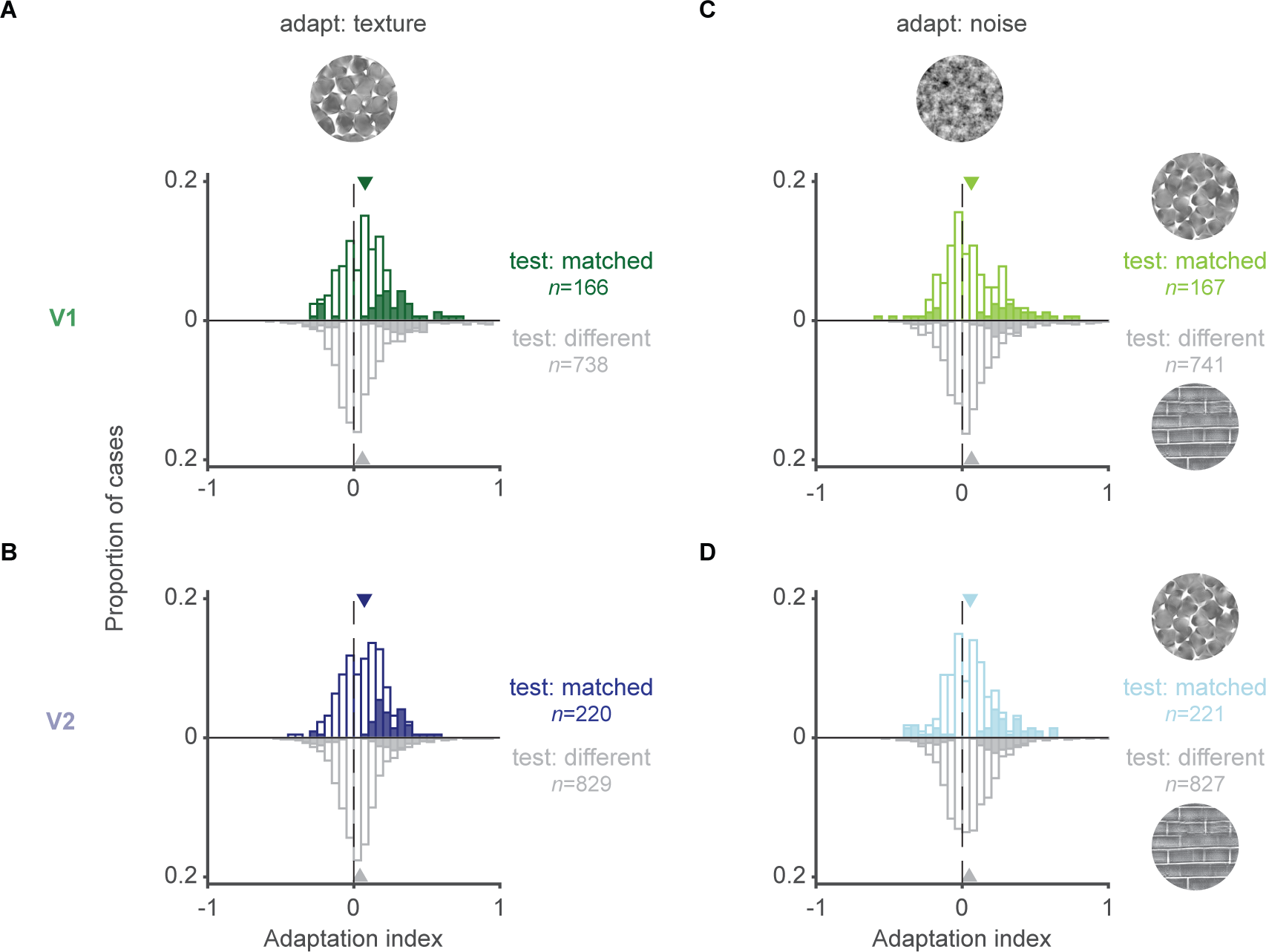
Adaptation effects between texture families. **(A, B)** Distributions of adaptation indices for V1(A) and V2 (B) units after adapting with a texture and testing with matched (colored traces, upright) or unmatched (gray, flipped) texture families. **(C, D)** Distributions of adaptation indices for V1 (C) and V2 units (D) after adapting with noise and testing with matched (colored traces, upright) or unmatched (gray, flipped) texture families. All adaptation indices are for test texture stimuli.

To test whether this adaptation specificity was due to the higher-order texture statistics or the spectral content of the image, we used a noise adapter and tested with textures matched and unmatched to the spectral content of the adapter. In V1, noise adapters induced similar effects for matched (0.06±0.01, p<0.001, t-test; Fig. 6C, upright) and unmatched (0.07±0.01, p<0.001, t-test; Fig. 6C, flipped) test texture families (p=0.53, unpaired t-test; Fig. 6C, top). In V2, noise adapters also induced effects of similar magnitude for matched (0.05±0.01, p<0.001, t-test; Fig. 6D, upright) and unmatched test textures (0.05±0.01, p<0.001, t-test; Fig. 6D, flipped; p=0.72, unpaired t-test for difference). These results indicate that the spectral differences between textures are not sufficient to induce adaptation specificity in V2 across texture families. Rather, adaptation specificity across texture families in V2 requires the presence of the higher-order statistics of different textures.

We wondered whether the degree of adaptation specificity for texture families was related to how well a given neuron differentiated among different textures. For instance, adaptation effects might be similar for two textures which evoke similar responses in a neuron, but different when one texture drove the cell more strongly than the other. To explore the relationship between texture selectivity and adaptation specificity, we focused on cases for which a single neuron’s responses allowed for reliable discrimination between the matched and a non-matched test texture (using a bootstrap analysis and requiring the discriminability to exceed the 95% confidence interval of the null distribution). We quantified single unit discriminability using the area under the receiver operating characteristic curve.

Texture families were slightly less discriminable in V1 (mean auROC: 0.14±0.004) than V2 (0.15±0.003, p=0.02, unpaired t-test), but the vast majority of V1 (88% of units) and V2 (94% of units) neurons responded differently to the matched test stimulus and at least one of the 4 non-matched textures. In the cases in which V1 units reliably discriminated between a texture pair (59.6% of cases), there was only slightly stronger adaptation indices for textures matched to the texture adapter (0.07±0.01) than for those from a different family (0.05±0.01, p=0.4, unpaired t-test). In V2, such cases (61.3% of cases) yielded clearly stronger adaptation indices for textures matched to the texture adapter (0.07±0.01) than for those that differed (0.01±0.01, p<0.0001, unpaired t-test).

In summary, we observe adaptation specificity in V2, but not V1, for texture families, meaning that responses to test textures matched to the adapter were reduced more strongly than responses to textures that were different from the adapter. This specificity required the presence of higher-level statistics in the adapter. The specificity was not strongly dependent on whether the individual unit responded differently to the various test textures.

### Effects of adaptation on selectivity and population information

Our analysis thus far compared changes in responsivity for different test stimuli, after adaptation with texture or noise stimuli. We next sought to understand how these changes in responsivity affected the information about textures in V1 and V2: does adaptation alter the selectivity of single neurons, or the information encoded by single neurons or populations? Do V1 and V2 neurons become better or worse at signaling differences between noise and texture images, or between different textures? The weaker responses after adaptation might be expected to reduce information, since the range of neuronal responses is compressed. On the other hand, the specificity of adaptation (when observed) might sharpen neuronal selectivity so that more information is encoded (Dayan and Abbott, 2001).

We first tested whether adaptation with a texture or noise stimulus altered the modulation index for test families matched to the adapter. We compared the modulation index, before and after adaptation, for units that were responsive to either texture or noise test stimuli before or after adaptation. Because the units that were responsive differed for texture and noise adapters, comparisons involved different units for the two adapters.

In V1, modulation indices for test stimuli matched to the adapter were unaffected by adaptation, for either texture (0.1±0.02 unadapted vs. 0.09±0.02 adapted, p=0.5 paired t-test; Fig. 7A) or noise adapters (0.1±0.018 vs. 0.1±0.02, p=0.9 paired t-test; Supplemental Fig. 1A). In V2, the modulation index was reduced slightly by both texture adapters (0.16±0.01 vs 0.13±0.01, p<0.001, paired t-test; Fig. 7B) and noise adapters (0.19±0.01 vs 0.17±0.01, p=0.002, paired t-test; Supplemental Fig. 1B). Thus, adaptation decreased how strongly neuronal responsivity in V2, but not V1, is modulated by higher-order texture statistics. A reduction in the modulation indices in V2 is expected given the stronger loss of responsivity for texture test stimuli than noise stimuli, after adaptation (Fig. 5).

**Fig. 7.**
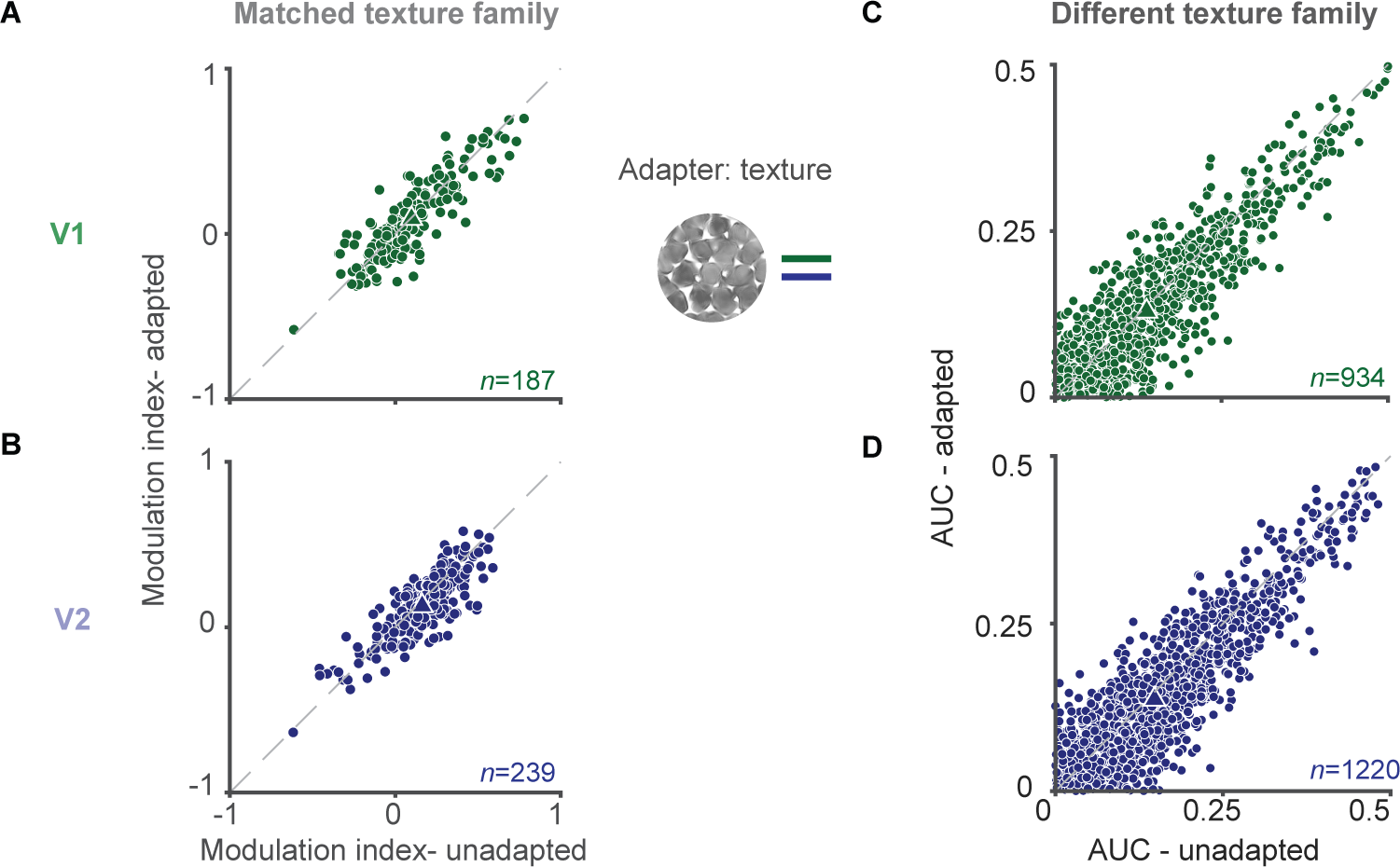
**(A, B)** V1 (A) and V2 (B) modulation indices for test textures matched to the adapter, before (abscissa) and after adaptation (ordinate) with a texture adapter. **(C, D)** V1 (C) and V2 (D) auROC values. Values are plotted before (abscissa) and after adaptation (ordinate) with a texture adapter.

We wondered if the change in modulation index in V2 was specific to the texture family of the adapter or if a similar loss of selectivity would occur for textures that were distinct from the adapter. We calculated modulation indices for the 4 texture families that were different from the adapter, for units that were responsive to either texture or noise of each family, before or after adaptation. In V1, there was no change in modulation indices after texture adaptation (0.01±0.006 unadapted vs. 0.02±0.006; p=0.24, paired t-test) but there was a slight reduction after adaptation with noise (0.01±0.006 vs. 0.002±0.006; p=0.01, paired t-test). In V2, modulation indices for texture families different from the adapter were not affected by adaptation, for either texture (0.05±0.006 vs 0.05±0.006; p=0.93, paired t-test) or noise adapters (0.05±0.006 vs. 0.05±0.006; p=0.53, paired t-test).

We next tested whether adaptation influenced discriminability between texture families. We quantified single unit discriminability using the area under the receiver operating characteristic curve (auROC). In V1, texture adapters slightly decreased discriminability (0.137±0.003 before adaptation compared to 0.129±0.003 after, p<0.0001, paired t-test; Fig. 7C). Discriminability was unchanged after noise adapters (0.136±0.003 compared to 0.135±0.004, p=0.12, paired t-test; Supplemental Fig. 1C). In V2, discriminability slightly decreased after presentation of texture adapters (0.148±0.003 to 0.137±0.003, p<0.0001, paired t-test; Fig. 7D) but less so after noise adapters (0.147±0.003 to 0.140±0.003, p=0.1, unpaired t-test; Supplemental Fig. 1D).

Finally, we tested whether adaptation altered texture discriminability in V1 or V2 neuronal populations. Though population effects might be expected to be closely related to changes in the discriminability afforded by single neurons, the two measures can diverge. For instance, changes in neuronal ‘noise’ correlations can enhance or mitigate the information loss observed in single neurons (Averbeck et al., 2006; Kohn et al., 2016).

We measured texture information in V1 and V2 populations by training a linear classifier to discriminate textures matched to the adapter from textures different from the adapter based on the measured neuronal responses, before and after adaptation (Fig. 8A shows example performance curves before and after adaptation). Classifier performance decreased after texture adaptation, in both V1 (0.84±0.02 vs. 0.8±0.02, p<0.001, paired t-test; Fig. 8B, top) and V2 (0.88±0.01 vs. 0.86±0.01, p= 0.02, paired t-test; Fig. 8B, bottom). Noise adapters reduced performance in V1 (0.84±0.02 vs. 0.82±0.02, p=0.004, paired t-test) but not V2 (0.88±0.01 vs. 0.87±0.01, p=0.2, paired t-test).

**Fig. 8.**
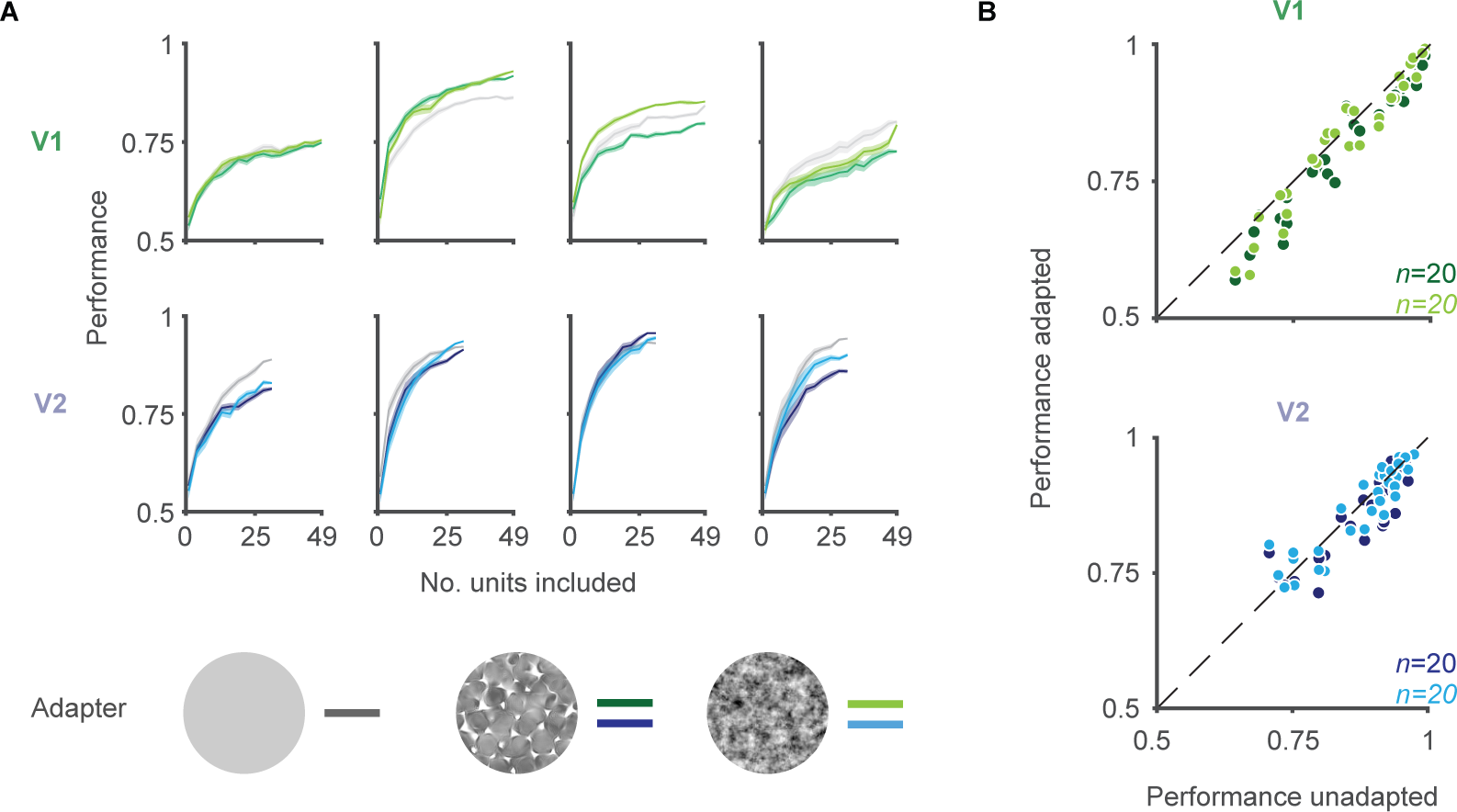
Classifier performance for texture discrimination. (**A**) Example performance curves for different texture family pairs (columns) in V1 (top, green) and V2 (bottom, blue). Discrimination was evaluated for units that were unadapted (gray traces) or adapted to a texture (dark green and blue) or noise stimulus (light green and blue). (**B**) Population sizes were matched across cases (n=30 neurons) to evaluate population performance before (abscissa) and after adaptation (ordinate) to textures (dark colors) and noise (light colors).

In summary, in both V1 and V2 adaptation either reduced information about textures (distinguishing between different textures) or texture statistics (distinguishing between textures and noise counterparts) or had no effect. Notably, in none of our comparisons did we observe that adaptation was helpful for visual encoding.

## Discussion

We measured neuronal adaptation in V1 and V2 to synthetic, naturalistic textures and their counterpart noise images. We found evidence for adaptation specificity for texture statistics in V2 but not V1, consistent with the different sensitivity of the two areas for these statistics. Though adaptation effects displayed some stimulus specificity, we found discriminability between textures and noise or between different texture families was either unaffected or impaired after adaptation.

### Adaptation to complex stimuli across stages of visual processing

Because visual processing occurs across stages of a hierarchical network, it can be difficult to determine whether adaptation effects observed in a particular area are generated there or reflect adapted input from other areas (i.e., inheritance). Previous work has attempted to infer the locus of neuronal adaptation by determining whether adaptation effects are specific for a stimulus feature for which selectivity first arises in the targeted area. For instance, these features might include sensitivity to stimulus orientation (Dragoi et al., 2000; Patterson et al., 2013), binocularity (i.e., testing whether adaptation transfer inter-ocularly; Sclar et al., 1985; Marlin et al., 1991), or spatial position (testing whether adaptation effects transfer across spatial locations within the receptive field of a targeted neuron; Marlin et al., 1991; Brown and Masland, 2001; Priebe et al., 2002; Kohn and Movshon, 2003; Dhruv and Carandini, 2014).

Using similar rationale, we tested whether adaptation with textures had different specificity in V1 and V2, consistent with the different sensitivity to texture statistics in those two areas. We observed differences between V1 and V2 effects, in a manner consistent with their differing selectivity.

First, the loss of V1 responsivity after adaptation with textures or noise images was statistically indistinguishable, for both texture and noise test images. This finding is consistent with V1 being ‘blind’ to the texture statistics: a texture and its noise counterpart are equivalent stimuli for most V1 neurons (Freeman et al., 2013; Ziemba et al., 2019) so no adaptation specificity would be expected. In contrast, in V2, adaptation effects depended on the presence of higher-order texture statistics. Texture adapters reduced responses to test textures more than noise images; a difference that was not as evident after noise adapters.

We also observed greater adaptation specificity in V2 than V1 across textures: V2 effects were strongest when adapter and test textures were from the same family. Surprisingly, there was little across-family specificity evident in V1. Though V1 neurons are largely insensitive to higher-order texture statistics, the different texture families also differed in spectral content. Since V1 adaptation effects are specific for spatial frequency (Movshon and Lennie, 1979; Saul and Cynader, 1989; Sharpee et al., 2006) and orientation (Nelson, 1991; Dragoi et al., 2000; Patterson et al., 2013), we might expect stronger V1 adaptation effects for matched versus non-matched textures. That this was not the case suggests that the spectral content of our textures were less different from each other than the specificity of the adaptation effects for these attributes. In V2, in contrast, neurons are sensitive both to the low-level (spectral) and higher-order statistics. Presumably the presence of additional stimulus features for which V2 neurons are sensitive could explain the specificity of adaptation effects across textures in that area.

Though we observed different adaptation specificity in V1 and V2, this specificity did not depend on the selectivity of the individual, sampled neuron in either area. Adaptation effects in V1 units with a high modulation index were still similar for textures and noise stimuli—that is, there was no specificity. Similarly, adaptation was not more specific for texture and noise stimuli in V2 units with a high modulation index. Thus, our results suggest that adaptation specificity reflects the selectivity of the local network rather than that of the sampled neuron per se. Previous studies have also found a lack of correlation between adaptation strength and tuning: in IT cortex, single-unit tuning for objects does not correlate with adaptation specificity (Sawamura et al., 2006); and in V1 adaptation effects are orientation specific even in cells with little or no orientation tuning (Wissig and Kohn, 2011).

Though a comparison of adaptation specificity in V1 and V2 neurons informs the locus of adaptation to textures, we note that the adaptation specificity we observe in V2 could occur because of how adaptation effects induced in V1 affect computations performed in V2 (see Patterson et al., 2014 for a discussion of how inherited adaptation can affect computations in higher visual areas). That is, it is possible that the changes in V1 responsivity give rise to specificity in V2 because of how V1 inputs to V2 are combined there. More specific statements of how adaptation effects arise in these two networks would require a formal model, describing how V1 inputs are combined to generate sensitivity to texture statistics in V2 and where in this model the plasticity induced by adaptation occurs.

### Methodological considerations

The loss of responsivity that we observed after adaptation was modest: adaptation indices were ∼0.1, reflecting a ∼20% reduction in response magnitude. Though this modest effect might be attributed to the texture and noise stimuli we used (which might drive neurons in early cortex less effectively than high contrast, sinusoidal gratings), prior work suggests that the duration of our adapters (0.6 s) is the more likely explanation. For instance, Patterson et al. (2013) found that V1 neuronal responses were reduced by ∼20% after 0.4 s adaptation with a high contrast, sinusoidal grating. Other studies using brief adapters in V1 have observed changes in responsivity of similar magnitude (Müller et al., 1999; Dragoi et al., 2002). Importantly, prior work suggests that more prolonged adaptation induces stronger effects than brief adaptation, but with similar specificity (Saul and Cynader, 1989; Dragoi et al., 2000; but see Patterson et al., 2013 for some exceptions). Thus, we would expect that prolonged adaptation with textures and noise would result in stronger effects than those reported here, but with similar specificity. Of course, effects of brief adaptation are most relevant for understanding how the visual system adjusts during natural experience, since saccades change the visual input within the receptive fields of neurons in early visual cortex every few hundred milliseconds (Yarbus, 1967).

Our experiments explored the effects of two different adapters on multiple test stimuli, each of which was presented many times to allow for accurate assessment of adaptation effects, including changes in population information. As a result, our measurements required stable recordings for long periods, which we accomplished by recording in sufentanil anesthetized animals. Unlike non-opioid anesthetics, sufentanil has little effect on cortical activity (Constantinople and Bruno, 2011). Additionally, previous work has shown adaptation effects to be similar in awake and anesthetized primates, including in V1 and area MT (Müller et al., 1999; Dragoi et al., 2000, 2002; Patterson et al., 2013, 2014b). Texture selectivity have been shown to develop gradually between V1 and V4, in a similar manner in anesthetized (Freeman et al., 2013; Ziemba et al., 2016, 2019) and awake primates (Okazawa et al., 2015, 2017). Thus, though we cannot exclude the possibility that our results were influenced by anesthesia, there is good reason to believe any such influence is weak.

Previous studies have found that stimulus size can influence adaptation effects because the suppressive influence of the surrounds adapts (Webb et al., 2005; Camp et al., 2009; Wissig and Kohn, 2012; Patterson et al., 2014a). Our stimuli were larger than the receptive fields of the sampled V1 and V2 neurons. Matching the stimuli to each neuron’s optimal RF size could produce different results. However, decreasing the stimulus size would also influence the texture statistics which are defined across spatial scales (Ziemba and Simoncelli, 2021). Because surround suppression for textures have been shown to be weaker than for gratings in V2 (Ziemba et al., 2018), stimulus size might influence adaptation less strongly than in previous gratings-based work (Wissig and Kohn, 2012; Patterson et al., 2013). Similarly, our approach of recording neurons whose receptive fields were not precisely centered on the texture may have led us to underestimate the sensitivity to texture statistics. However, it is unlikely that this led to an appreciable underestimate of adaptation specificity for those statistics, as we did not observe a relationship between the adaptation specificity and the degree of sensitivity for texture statistics in our recorded sample.

### How neuronal selectivity is altered by adaptation

Adaptation effects are often proposed to allow sensory systems to better encode stimuli similar to those recently encountered—a ‘fine-tuning’ of the system to current environmental demands. Light adaptation in the retina clearly does this, allowing us to discriminate between luminance values that vary over many orders of magnitude with receptors that have a more limited dynamic range (Shapley and Enroth-Cugell, 1984; Rieke and Rudd, 2009). However, it has been difficult to establish the benefit of many cortical neuronal adaptation effects (Kohn, 2007; Solomon and Kohn, 2014; Webster, 2015).

We assessed whether adaptation altered texture encoding and found, by several measures, that it worsened it. First, adaptation reduced the magnitude of the modulation index in V2, meaning that single neurons responded more similarly to textures and noise stimuli after adaptation, because responses to textures were reduced more strongly than those to noise stimuli. The reduction in the modulation index was evident only in textures matched to the adapter. In V1, where modulation indices were smaller, there was no evident change in the modulation index after adaptation.

Second, V1 and V2 responses afforded weaker discriminability between different textures. This was evident as a slight loss in the classifier performance in discriminating responses to different textures, using the measured population responses. In addition, single V2 neurons responded less distinctly to different textures after adaptation, which was evident as a decrease in single neuron texture discriminability.

In none of our analyses did we find that adaptation improved performance for discriminating textures. Some previous studies have found improved neuronal discriminability after adaptation, for simple stimulus features like orientation or contrast (Müller et al., 1999; Gutnisky and Dragoi, 2008; Jin et al., 2019 but see Dragoi et al., 2002; Ghodrati et al., 2019 for counterinstances). Our results are more consistent with psychophysical studies in humans and non-human primates, which find that discrimination is impaired when the adapter is similar to the target stimuli (Dragoi et al., 2002; see Clifford, 2002 for review). Alternatively, our results might suggest that improvements in neuronal discriminability for simple stimulus features might not be evident for more complex stimuli like naturalistic textures.

## Supplementary Figure

**Supplemental Fig. 1.**
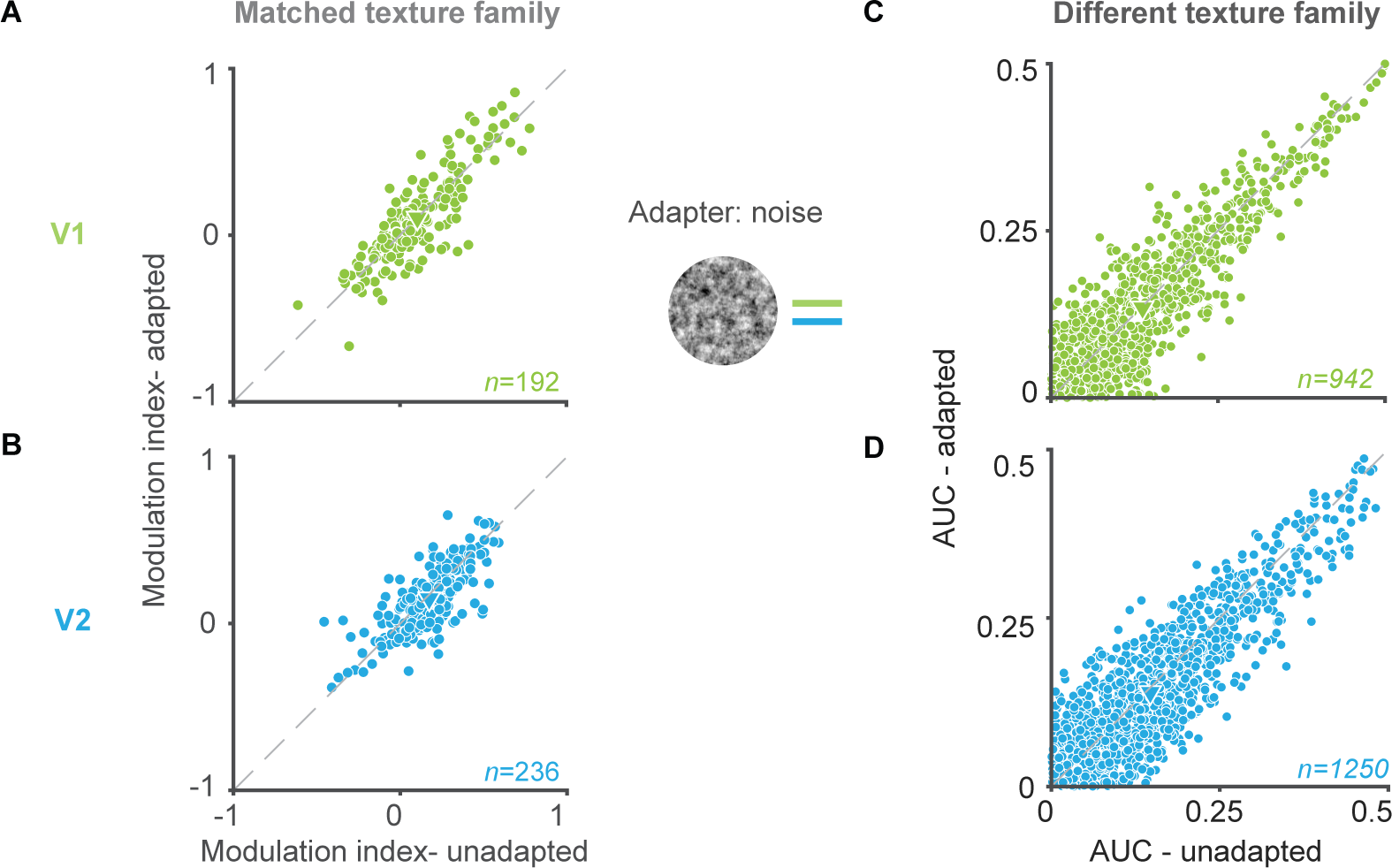
**(A, B)** V1 (A) and V2 (B) modulation indices for test textures matched to the adapter, before (abscissa) and after adaptation (ordinate) with a noise adapter. **(C, D)** V1 (C) and V2 (D) auROC values. Values are plotted before (abscissa) and after adaptation (ordinate) with a noise adapter.

